# For an early and strong immune response in the monomerization of polymeric immunoglobulins intermonomeric interaction relationship of potassium hydroxide

**DOI:** 10.1101/2022.11.13.516299

**Authors:** Emin Zümrütdal

## Abstract

Immunoglobulins (Ig), which are an integral part of the immune system in humans and animals, continue to surprise people with their effectiveness and importance every day since they were discovered for 2 centuries.

Although the avidity of polymeric immunoglobulins (pIg) in the lumen in the respiratory system is high, its mobility is limited and unlike monomeric immunoglobulins (Ig), it can carry the antigenic (Ag) structure to the lamina propria by active transport. pIg’s have a high weight because they have a large number of monomers. Disulfide bonds are important in the bonds that connect Ig monomers, and disulfide bonds have critical functions in construction, both in the J chain and in the intermonomeric domain. Disulfide bonds can interact quite easily with hydroxyl(OH^−^) ions. It is a strong molecular structure that gives OH^−^ ions in potassium hydroxide (KOH).

In our study, the potential of pIg to be made monomeric was investigated in order to increase the mobilization, antigen affinity and antigen avidity of pIgs (IgM and IgA) in the secretion in the lumen.

For this purpose, intermonomeric disulfide bonds were determined by investigating cysteine locations in the J chain and on the monomers. In this study, by targeting these disulfide bonds, the intermolecular interaction energies with KOH for the destruction of these bonds were evaluated by in silico studies.

Exergonic intermolecular free energy interactions were detected between the disulfide bonds in the J chain and in the intermonomeric domain and the KOH molecule.

**Graphical abstract:** Graphic with abstract of the study.

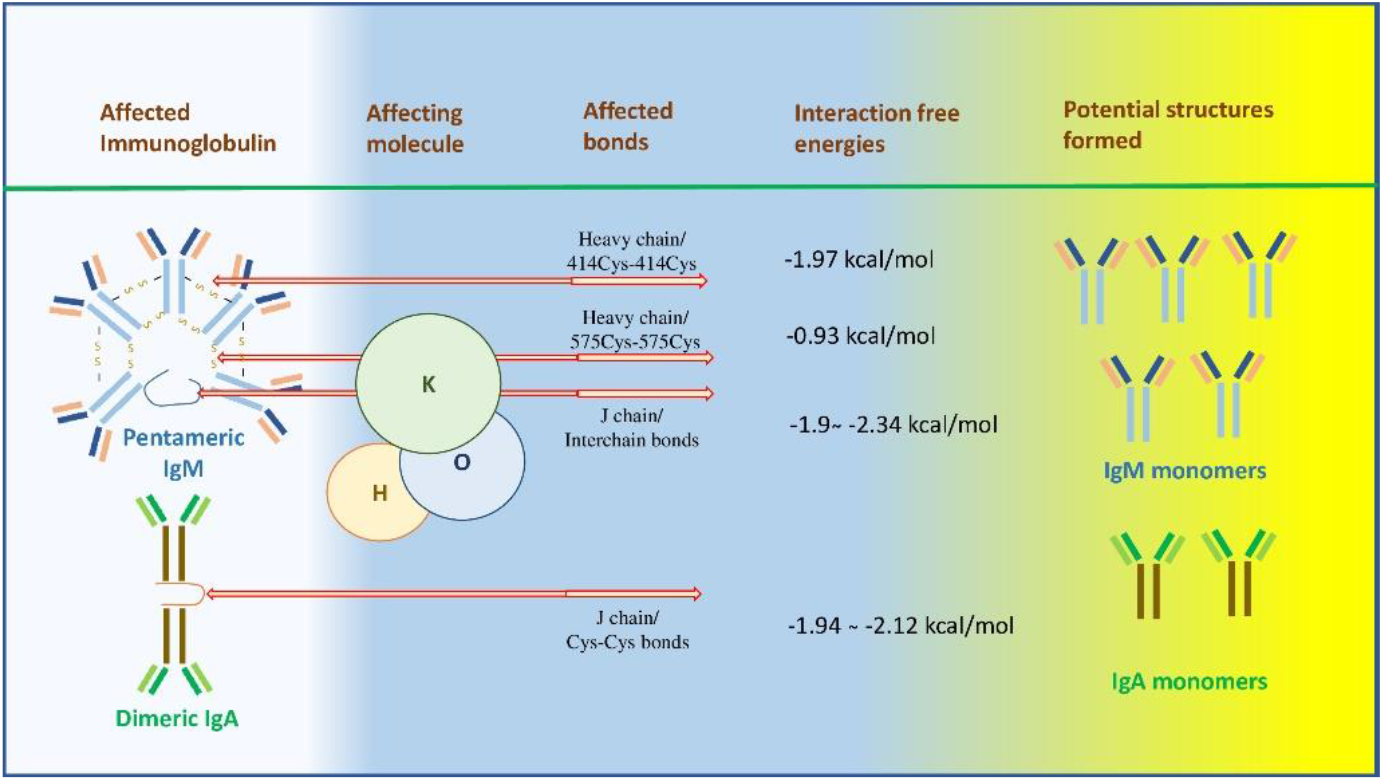

## Introduction

Immunoglobulins (Ig) were defined by scientists at the end of the 19th century and their magnificent role in the immune response was revealed. As the importance of these molecules produced by B lymphocytes in body defense became clear, the interest in Ig’s increased even more. At the end of the 20th century, the discovery of Ig, which has a different structure but functionality in animals such as sharks and llamas, gave rise to new hopes in the treatment of various diseases(1). The positive efficacy of the different Ig structures discovered in the treatment and the increasing technological developments have led scientists to the discovery of Ig or Ig fragments with different structures (2). However, despite technological advances and knowledge, the therapeutic index of Ig produced has been a controversial problem(3). In addition, the high cost of treatment makes it difficult to access these treatments.

In the host defense system, Ig’s capture molecular structures that are not compatible with body cells, such as viruses, bacteria or cancer cells, and present them to B lymphocytes, macrophages or dendritic cells, or they stimulate chemotaxis by opsonizing. This functionality demonstrates the importance given to Igs very well. After the antigen (Ag) molecules are captured, one of the places of presentation to the phagocytic cell or lymphocyte is the lamina propria under the mucosa in the respiratory lumen(4). If Ag molecules coming to the lamina propria are not captured and eliminated, they spread throughout the body and cause various diseases.

Mucous surfaces in the respiratory lumen encounter Ag molecules very frequently. In the first line of defense, hair, mucus, ciliary activity and epithelium, which are involved in innate immunity, have an important place (5). In the respiratory lumen, the Ig’s waiting in the secretion on the mucosa act as the soldiers of the defense system and prevent the Ag molecules that come up to the mucosal surface from freely passing into the lamina propria. IgA is the most released Ig, which forms the defense line in the re The transport properties of Ig from the basolateral area in the respiratory lumen to the apical region, which is the line of defense, differ according to the monomer number of Ig.

While IgG, IgE and IgD have a monomeric structure, IgA (dimer) and IgM (pentamer & hexamer) are immunoglobulins (pIg) with a polymeric structure. IgA, which has a dimeric structure, forms pIg by linking 2 monomers with the J chain in the Fc part. In the IgM pentamer structure, in addition to the disulfide bonds in the J chain, it is held together by both the Fc part and intermonomeric disulfide bridges (7).spiratory lumen, followed by IgM, and small amounts of other Ig may be found (6,7).

Mucosal transport of monomeric Ig’s can be by paracellular passive transfer, whereas pIg’s must be bound by special molecular structures for their transfer to the mucosal surface(8). In addition to forming the pIg structure, the J chain also interacts with the pIg Receptor/Membrane Secretory Component (pIgR/SC) and has an important role in the active transport of pIg from the basolateral region to the apical region(6,9).

If Ag is present on the apical surface, it is identified and captured by Ig and brought to the basolateral part. Here, Ag captured by Ig is presented to B lymphocytes and phagocytic cells. Stimulation of B lymphocytes and chemotaxis are activated and body defense begins.

However, while increasing monomeric structures in pIg in these functions increase avidity as an advantage, active transport is required in paracellular transport as a disadvantage. This limits the mobility of plg’s.

Disulfide bonds play an important role in the construction of pigs and the J chain. However, when disulfide bonds encounter alkaline molecular structures, they can change the bond structure by forming stable compounds(10-12). Potassium hydroxide (KOH) is one of these alkaline molecular structures and can make nucleophilic attacks(13).

Acidity increases in tissues with inflammation. The increase in acidity and inflammation may result in new Ag loads due to tissue damage caused by oxidative stress. KOH can have a different positive effect by blocking the formation or increase of acidity with the OH^−^ ions it contains (14).

In the light of this information, the intermolecular interactions of disulfide bonds, one of the structures that play a binding role in the skeletal structure of pIg’s, with KOH were evaluated with in silico (Docking) studies created with artificial intelligence.

## Methods

### Receptor and ligand retrieval

Structure of dimeric sIgA complex (IgA, PDB ID: 6UE7, resolution 2.90 Å), and the cryo-EM structure of human IgM (IgM-Fc, PDB ID: 6KXS, resolution 3.40 Å) in complex with the J chain were downloaded from the RCSB PDB database (https://www.rcsb.org/) and were used as target receptors in the molecular docking study. The KOH simulated as the reacting ligand against these human Igs was drawn using GausView 6.0.16 embedded in Gaussian 09 program and optimized with the B3LYP hybrid function (6-311G (d,p) basis set) (15).

Therefore, in molecular docking studies with KOH, Cys13-Cys101, Cys72-Cys92, Cys109-Cys134 S-S bonds were targeted on the dimeric IgA (PDB ID: 6UE7) molecule. On the other hand, in the IgM pentameric (PDB ID: 6KXS) structure, Cys12-Cys100, Cys108-Cys133, Cys14-Cys575 and Cys68-Cys575 on the J chain were targeted, while on the F and G chain, Cys414-Cys414, Cys575-Cys575 and Cys474-Cys536 S-S bonds were selected as target regions that could be attacked by KOH.

### Molecular docking between KOH and human Ig structures

Prior to docking simulations with AutoDock Vina, target (IgA and IgM) and ligand (KOH) structures were prepared with AutoDockTools 1.5.6 and saved in pdbqt format (16,17). In molecular docking simulations with human Igs and KOH, polar hydrogen atoms in receptors and ligand molecules were retained, whereas, non-polar hydrogens were merged. Kollmann charges were assigned to the receptors and Gasteiger charges were assigned to the KOH.

Targeted disulfide bond locations in both Ig molecules prior to molecular docking are shown in **Figure 1** and **Figure 2**.

**Figure 1.**
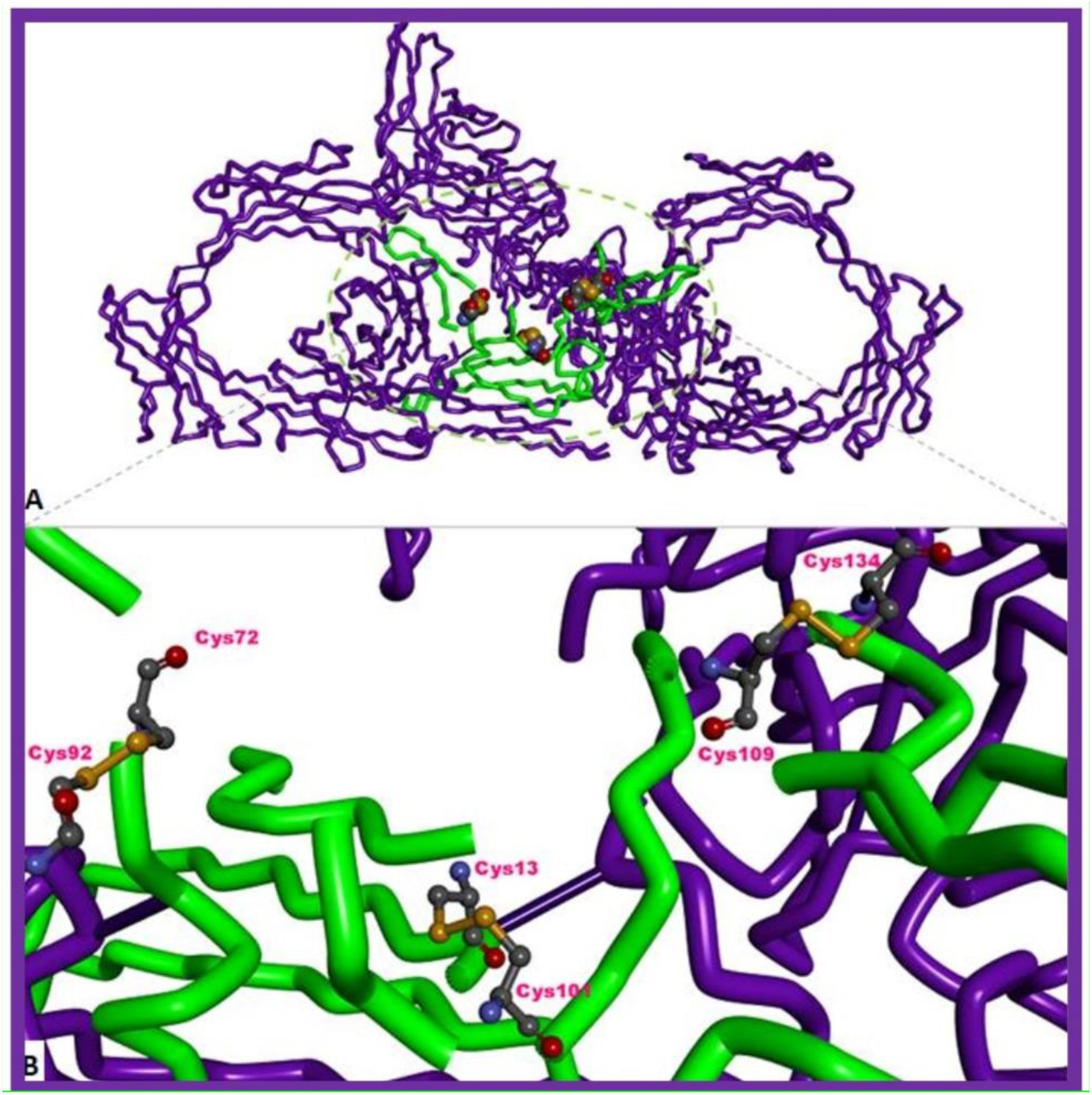
3D structure of human sIgA, (DPB ID: 6UE7) and its targeted region in molecular docking simulations. **A**. 3D tube view of dimeric slgA and the targeted J-chain (in green color surrounded by green dashed lines) along with the CPK view of amino acid residues involved in disulfide bonding. **B**. Zoom-in view of targeted amino acid residues and disulfide bonds (in yellow) on the J-chain prior to docking. Residues are given in the ball and stick mode. Image rendering was performed using DS Studio Visualizer v16.

**Figure 2.**
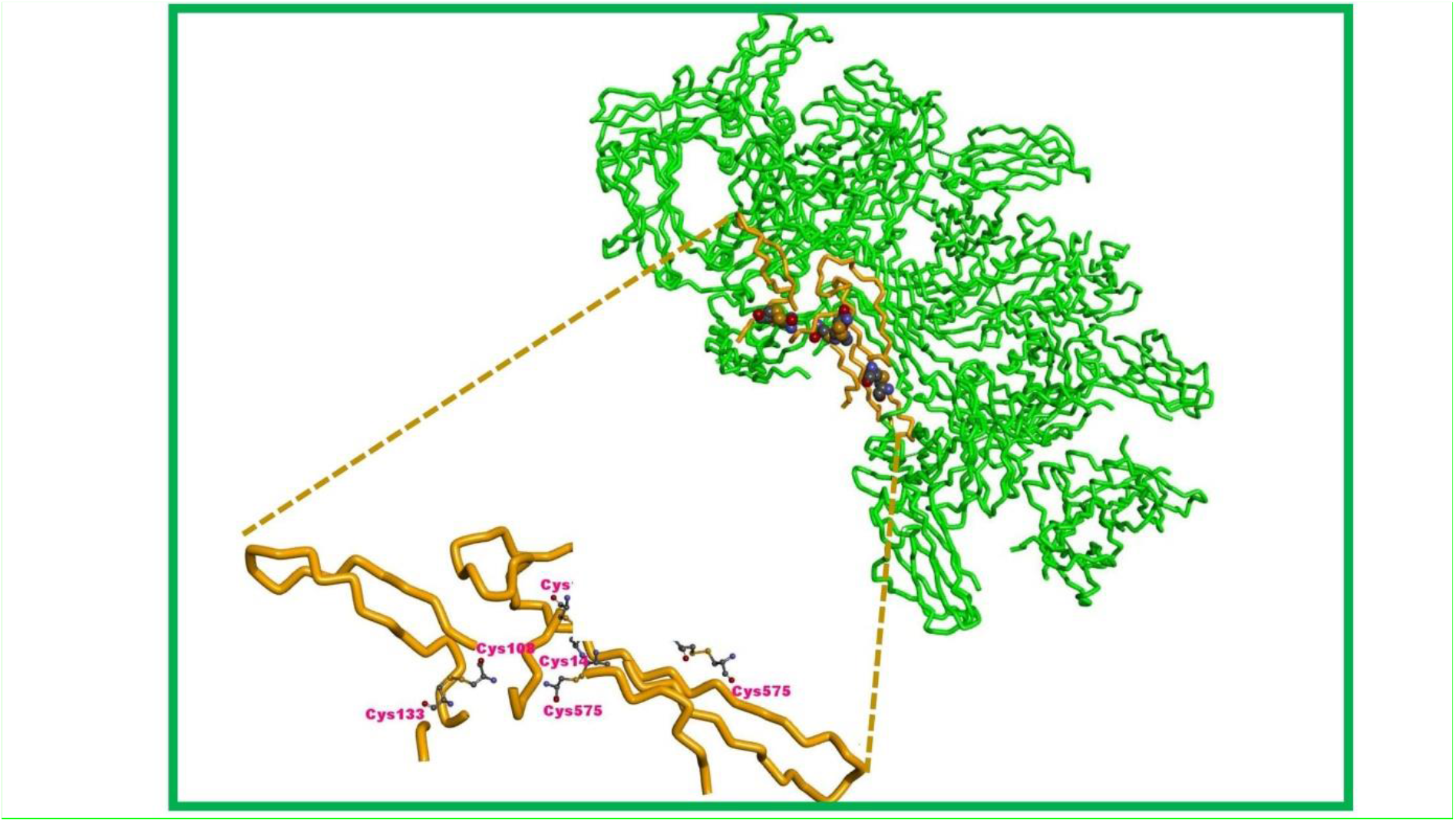
**A**. 3D overview of human IgM pentameric structure and its targeted regions in molecular docking simulations. **B**. Zoom-in 3D tube view of J-chain (in orange color surrounded by black dashed lines) along with the ball and stick view of the targeted amino acid residues involved in disulfide bonding. Image rendering was performed using DS Studio Visualizer v16.

On the J chain of the dimeric IgA molecule, the following grid box sizes are set for the Cys108-Cys133, Cys14-Cys575, Cys12-Cys100 and Cys68-Cys575 S-S bonds, respectively: 10×10×10 Å points (x = 154.13; y = 148.47; z=170.24); 10×10×10 Å points (x = 144.67; y = 138.62; z = 160.57); 10×10×10 Å points (x = 152.19; y = 137.11; z = 159.81) and 10×10×10 Å points (x = 146.01; y = 122.12; z = 160.78). In docking calculations against the J chain of IgM, the following grid box sizes were set: 10×10×10 Å points (x = 154.13; y = 148.47; z = 170.24) for Cys108-Cys133 S-S bond; 10×10×10 Å points (x = 144.67; y = 138.62; z = 160.57) for Cys14-Cys575 S-S bond; 10×10×10 Å points (x = 152.19; y = 137.11; z = 159.81) for Cys12-Cys100 S-S bond; and 10×10×10 Å points (x = 146.01; y = 122.12; z = 160.78) for Cys68-Cys575 S-S bond. For the two different S-S bonds on the F and G chain of IgM, the following grid box sizes are set for the docking: 7×7×7 Å points (x = 105.45; y = 99.15; z = 94.13) for the Cys414-Cys414 S-S bond on the F chain, a grid box size of 7×7×7 Å for the Cys575-Cys575 S-S bond on the F chain, and a grid box size of 10×10×10 Å points (x = 122.89; y = 103.99; z = 118.03) was set for the Cys474-Cys536 S-S bond in the G chain.

In this study, the close vicinity of a total of 10 S-S disulfide bonds with which KOH can interact was selected as target regions in docking simulations.

After ten separate docking runs against a total of three target regions on IgA and IgM (IgA J chain, and IgM J, F and G chains; 20 dockings for each S-S bond) of KOH, all potential binding modes of this ligand were clustered by AutoDock Vina 1.2.0 and were ranked based on the binding free energy (ΔG°; kcal/mol) of the ligand conformation with the lowest binding free energy against each S-S bond described above. Since no interaction with a single KOH molecule was observed between the S-S bonds of Cys414-Cys414 amino acids in IgM, these bonds were specifically studied with multiple KOH ligands. The top-scored docking conformations of KOH calculated by AutoDock Vina 1.2.0 among different poses against the IgA and IgM receptors were rendered and analyzed for intermolecular interactions using Discovery Studio Visualizer v16 (18).

## Results

The intermolecular interactions of KOH in the vicinity of the Cys72-Cys92 disulfide bond in the J chain of the human dimeric IgA molecule are shown in supplement file. In this region, KOH entered into electrostatic and metal-acceptor interactions with the Asp73 residue via its potassium atom, however, in this docking pose, it was reasonably far to the Cys72-Cys92 S-S bond. KOH displayed an energetically favorable binding affinity with the Asp73 residue (ΔG°=-2.12 kcal/mol, **Table 1**).

**Table 1.**
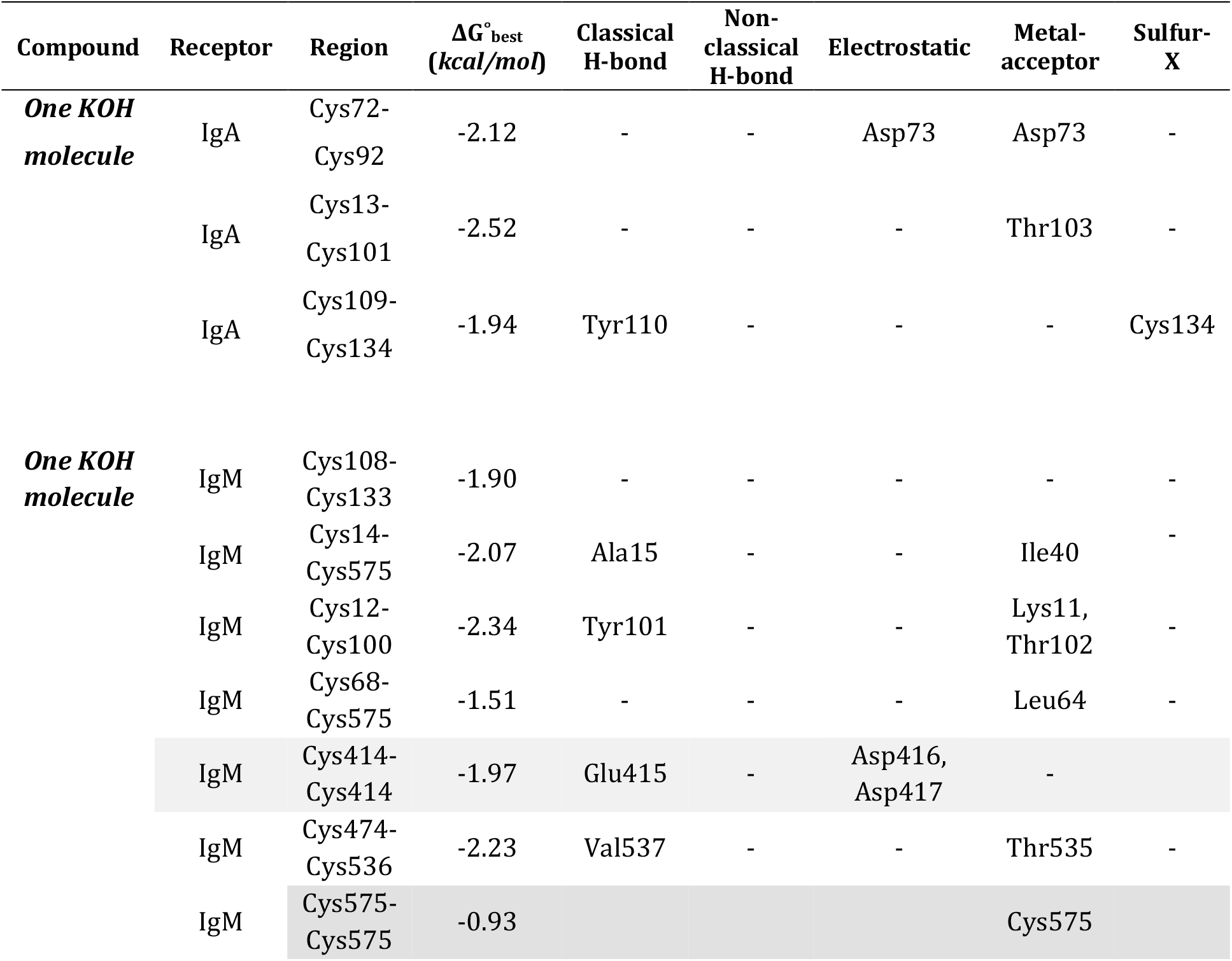
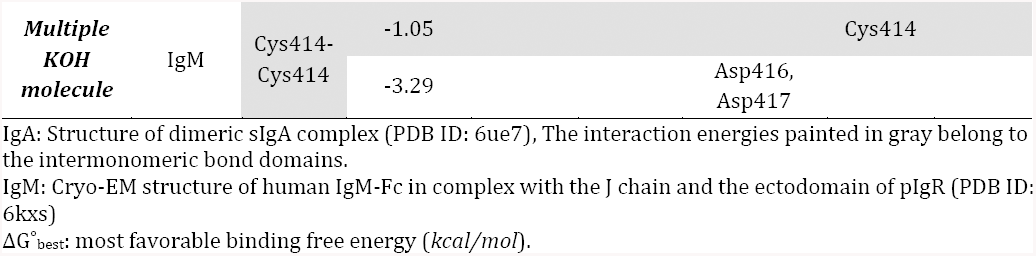
Intermolecular interactions of KOH in the vicinity of some disulfide bonds in human IgA (dimeric) and IgM (pentameric) molecules and the calculated binding free energies (ΔG°) of these interactions.

KOH formed a metal-acceptor interaction with Thr103 via its potassium atom near the Cys13-Cys101 disulfide bond in the J chain of dimeric IgA (The figure and its explanation are in the supplement file). The position of KOH in this region (its proximity to the disulfide bond) was more favorable than its position in the Cys72-Cys92 region, but interacted with a different amino acid (Thr103) than the amino acids involved in the disulfide bond (The figure and its explanation are in the supplement file). KOH showed an energetically favorable binding affinity (ΔG°=-2.52 kcal/mol) with the oxygen atom in the side chain of Thr103 near the Cys13-Cys101 domain (**Table 1**).

KOH, through its oxygen atom, formed a Sulfur-X (S-O) bond with the sulfur (S) atom of Cys134 in the Cys109-Cys134 S-S bond region of the J chain of dimeric IgA (**Figure 3**).

**Figure 3.**
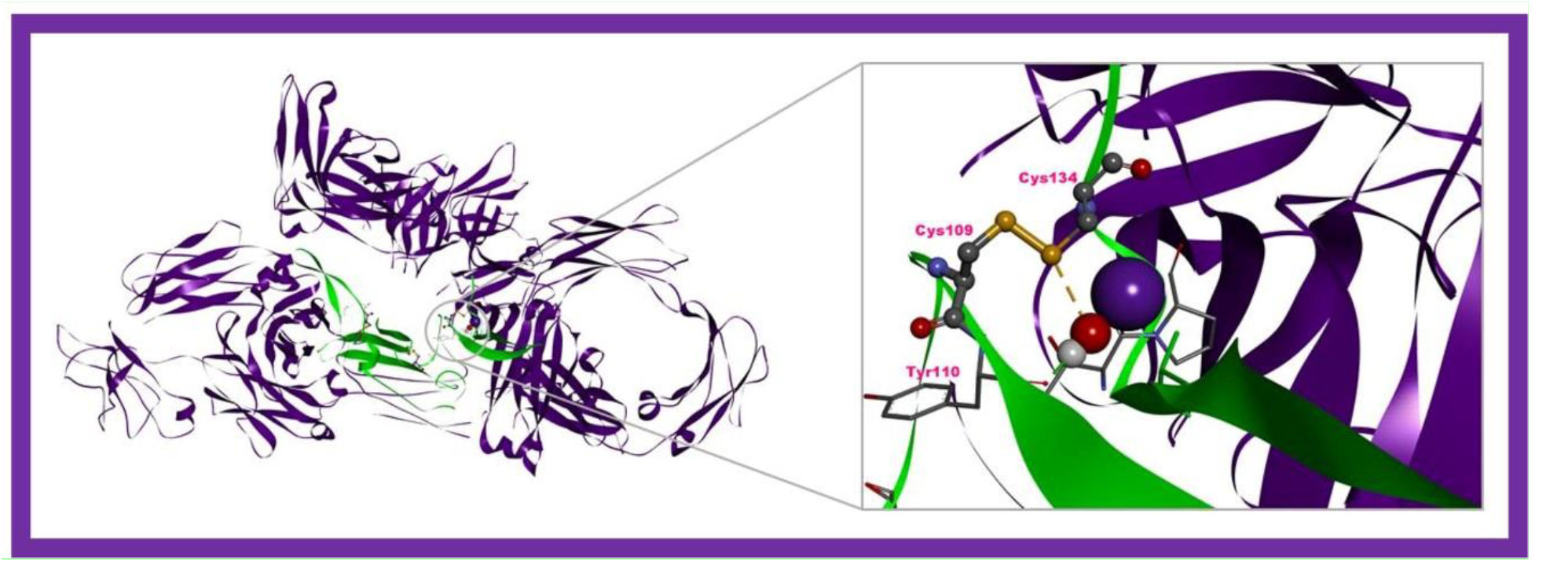
Top-ranked conformation of KOH in complex with the human sIgA in the vicinity of Cys109-Cys134 disulfide bond. The yellow dashed line represent electrostatic, whereas the green dashed line represent hydrogen bonding interactions, respectively. Image rendering was performed with DS Studio v16 software.

The binding free energy of this interaction was found to be chemically favorable (ΔG°=-1.94 kcal/mol, **Table 1**).

In the second part of our study, we simulated the affinity of KOH against Cys-Cys disulfide bonds located in two different regions (the J chain, and the F-G chain dimer) on the human IgM pentameric structure, and examined the resulting intermolecular interactions.

For this purpose, the affinities of KOH against Cys108-Cys133, Cys14-Cys575, Cys12-Cys100, Cys68-Cys575, Cys474-Cys536, Cys414-Cys414 and Cys575-Cys575 S-S bonds were calculated. The intermolecular interactions of KOH in the vicinity of the Cys108-Cys133 S-S bond in the J chain of IgM are given in **Figure 4A**.

**Figure 4.**
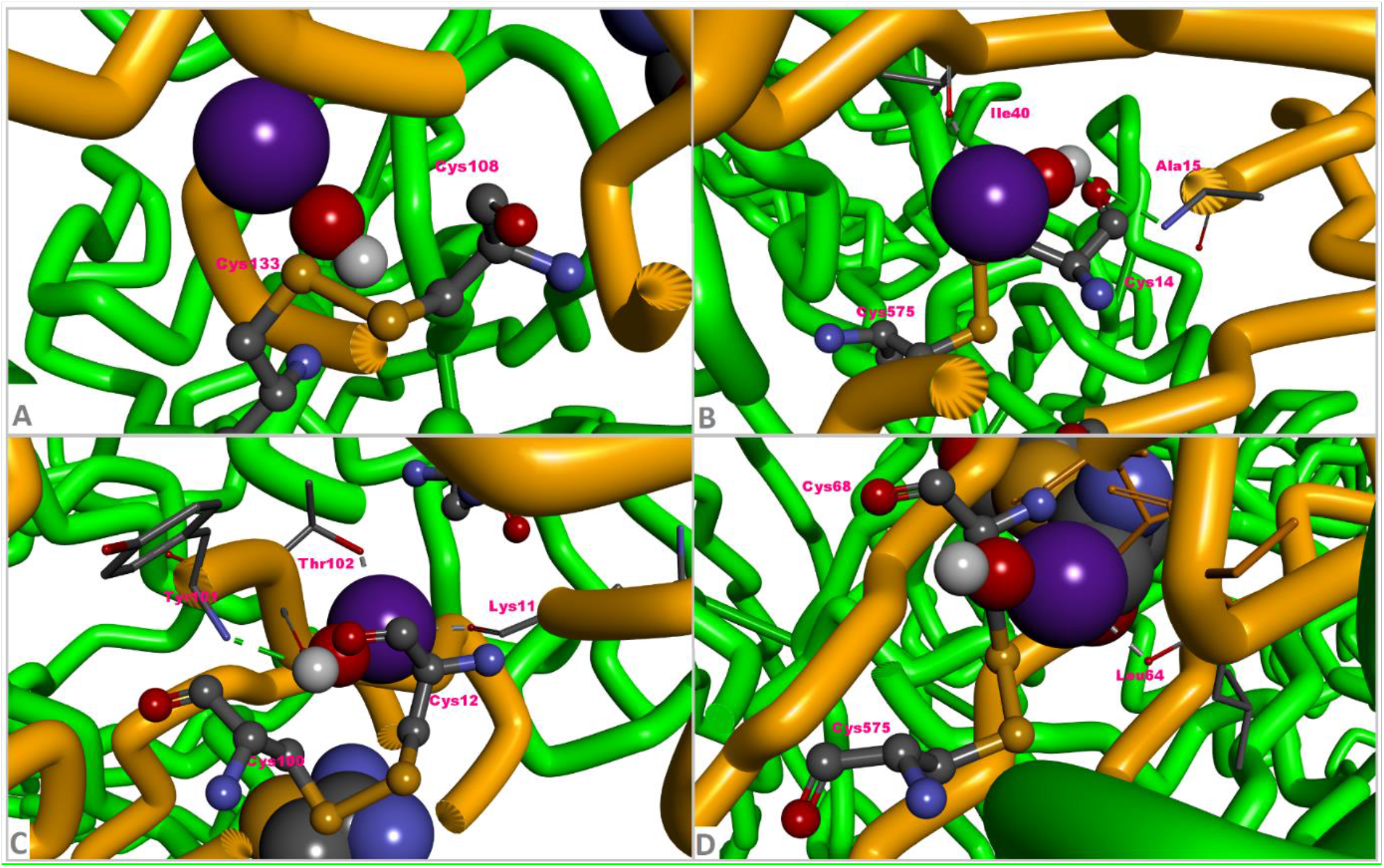
Intermolecular interactions of KOH near disulfide bonds in the J chain of human Ig M (IgM, PDB ID: 6KXS). **A**. KOH interaction in the vicinity of Cys108-Cys133 disulfide bond. **B**. KOH interaction in the vicinity of Cys14-Cys575 disulfide bond. **C**. KOH interaction in the vicinity of Cys12-Cys100 disulfide bond. **D**. KOH interaction in the vicinity of Cys68-Cys575 disulfide bond. Ig domains were depicted in the tube mode, and the aminoacids in interaction with KOH were given in the ball and stick mode. Image rendering was achieved with DS Studio Discovery Visualizer v16.

KOH did not interact with the Cys108-Cys133 S-S bond or different residues in this region, but the resulting reaction was energetically favorable (ΔG°= -1.90 kcal/mol, **Table 1**). Most likely, KOH has established weak Van der Waals interactions with amino acid side chains in its immediate vicinity in this region. Intermolecular interactions of KOH near the Cys14-Cys575 S-S disulfide bond are given in **Figure 4B**.

KOH formed a hydrogen (H) bond with the nitrogen (N) atom of Ala15 and metal-acceptor interaction with the oxygen (O) atom of Ile40 (**Figure 4B**). Although KOH was located very close to the Cys14-Cys575 S-S bond in this region, it showed no reaction against the S atoms participating in the bond. The affinity of KOH in this region was found to be energetically favorable (ΔG°= -2.07 kcal/mol, **Table 1**).

The intermolecular interaction of KOH around the Cys12-Cys100 S-S bond is given in **Figure 4C**. KOH formed an H-bond with Tyr101 just across the S-S disulfide bond, and a metal-acceptor bond both with Lys11 and Thr102 (**Figure 4C**). However, no interaction with the disulfide bond in this region (Cys12-Cys100) was occured. The affinity of KOH for IgM pentamer in this predetermined coordinate is energetically favorable (ΔG°= -2.34 kcal/mol, **Table 1**).

Intermolecular interactions of KOH around the Cys68-Cys574 S-S bond are given in **Figure 4D**. KOH formed a metal-acceptor interaction with Leu64 in this region and its binding affinity was found to be energetically favorable (ΔG°= -1.51 kcal/mol, **Table 1**).

The intermolecular interactions of KOH in the vicinity of the Cys414-Cys414 and Cys474-Cys536 S-S bonds in the F-G chain of IgM are given in **Figure 5**. KOH formed a H-bond with Glu415 near the Cys414-Cys414 disulfide bond and electrostatically interacted with Asp416 and Asp417 residues (**Figure 5A**). The binding affinity of KOH in this region of IgM is energetically favorable (ΔG°= -1.97 kcal/mol, **Table 1**). Intermolecular interactions of KOH in the Cys474-Cys536 S-S bond region of IgM are given in **Figure 5B**. In this region, KOH formed a H-bond with Val537 and a metal-acceptor interaction with Thr535 (**Figure 5B**). The binding affinity of KOH towards this region of IgM was also energetically favorable (ΔG°= -2.23 kcal/mol, **Table 1**).

**Figure 5.**
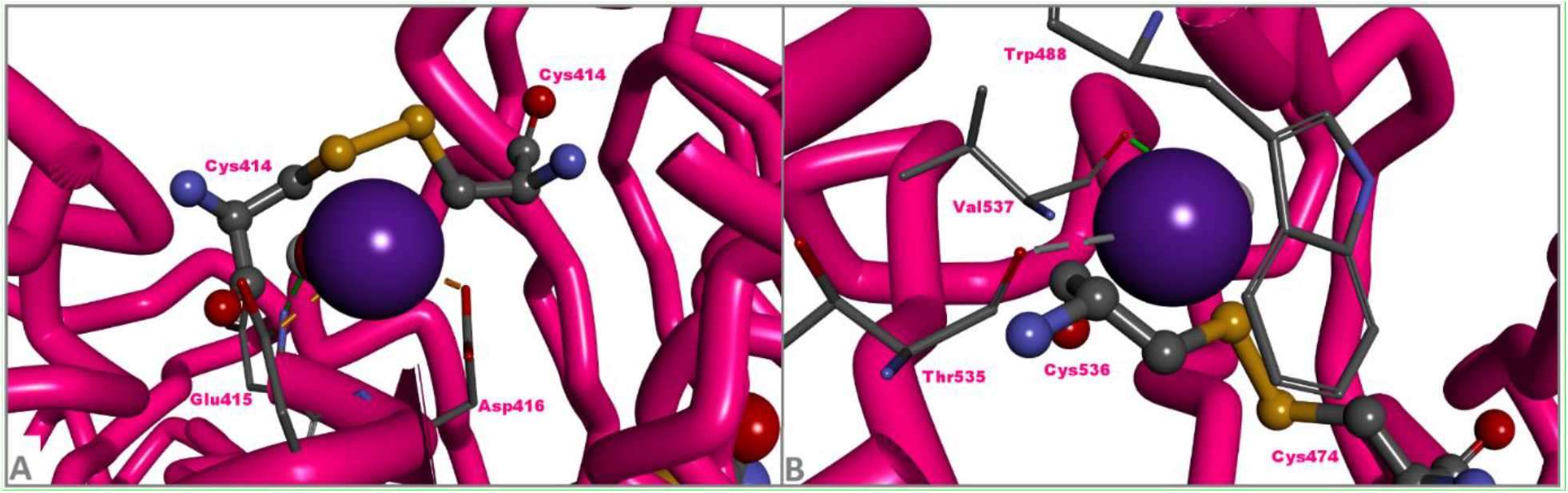
Intermolecular interactions of potassium hydroxide (KOH) near Cys-Cys disulfide bonds in the F and G chains (Fab and Fc regions) of human pentameric immunoglobulin M (IgM, PDB ID: 6KXS). **A**. KOH interaction in the vicinity of Cys414-Cys414 disulfide bond. **B**. KOH interaction in the vicinity of Cys474-Cys536 disulfide bond. Antibody domains were depicted in the tube mode, and the amino acid residues in interaction with KOH were given in the ball and stick mode. Image rendering was performed with DS Studio Discovery Visualizer v16.

However, when KOH molecules were increased in the medium, a interaction was detected between the KOH ligand and the sulfur atom of the cysteine amino acid(**Figure 6**).

**Figure 6:**
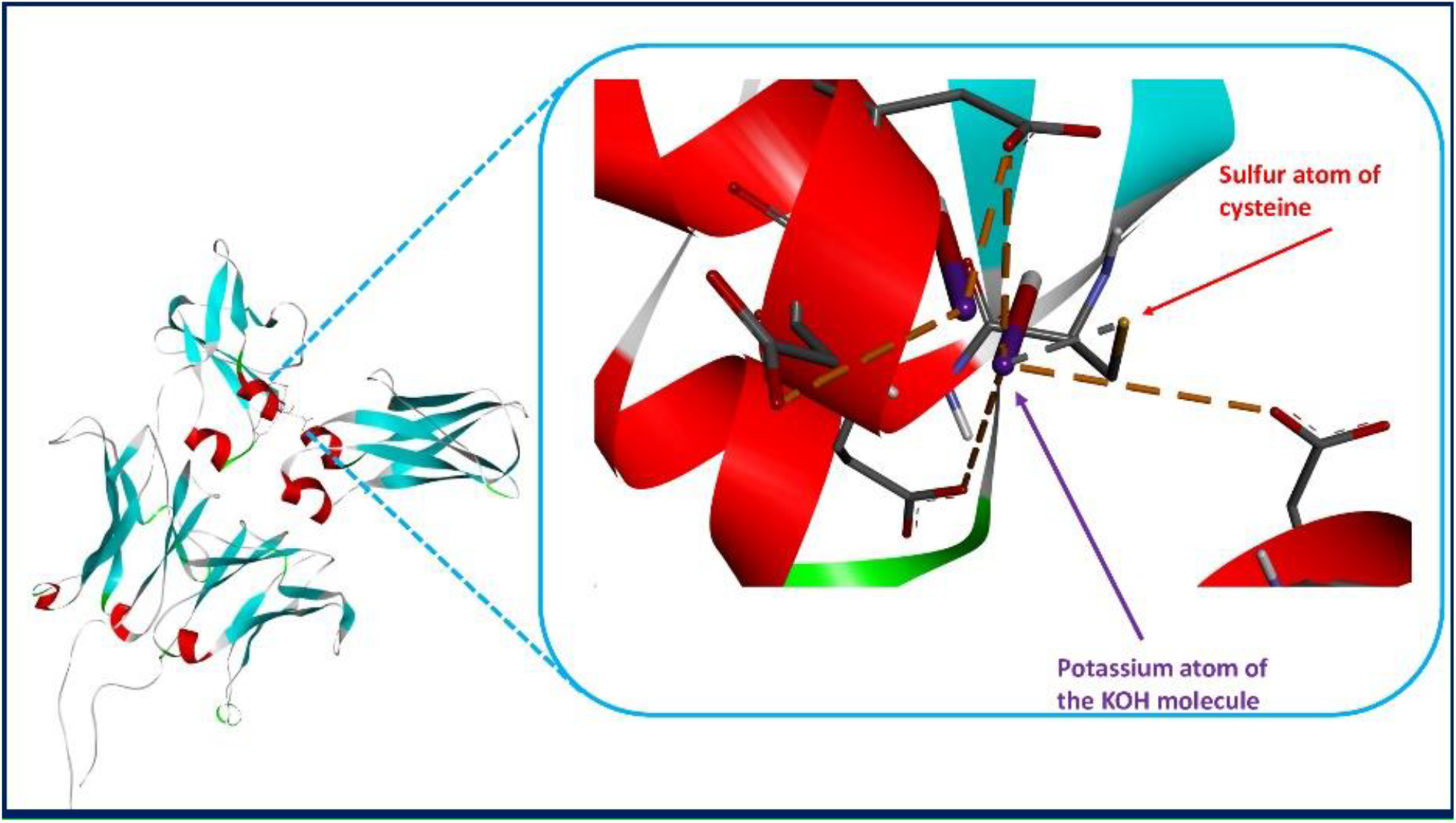
The relationship between the KOH ligand and the sulfur atom of the 414th cysteine amino acid in the IgM G chain and between other nearby amino acids.

In the simulation study with multiple KOH molecules, the electric charge of the oxygen atom to which the potassium atom bonded was -0.63e, while the electric charge of the sulfur atom was -0.09e.

In IgM, metal-acceptor interaction was detected between the potassium atom and the sulfur atom in the disulfide bond between cysteines, which are the 575th amino acids of the G and F chains(Figure 7).

**Figure 7:**
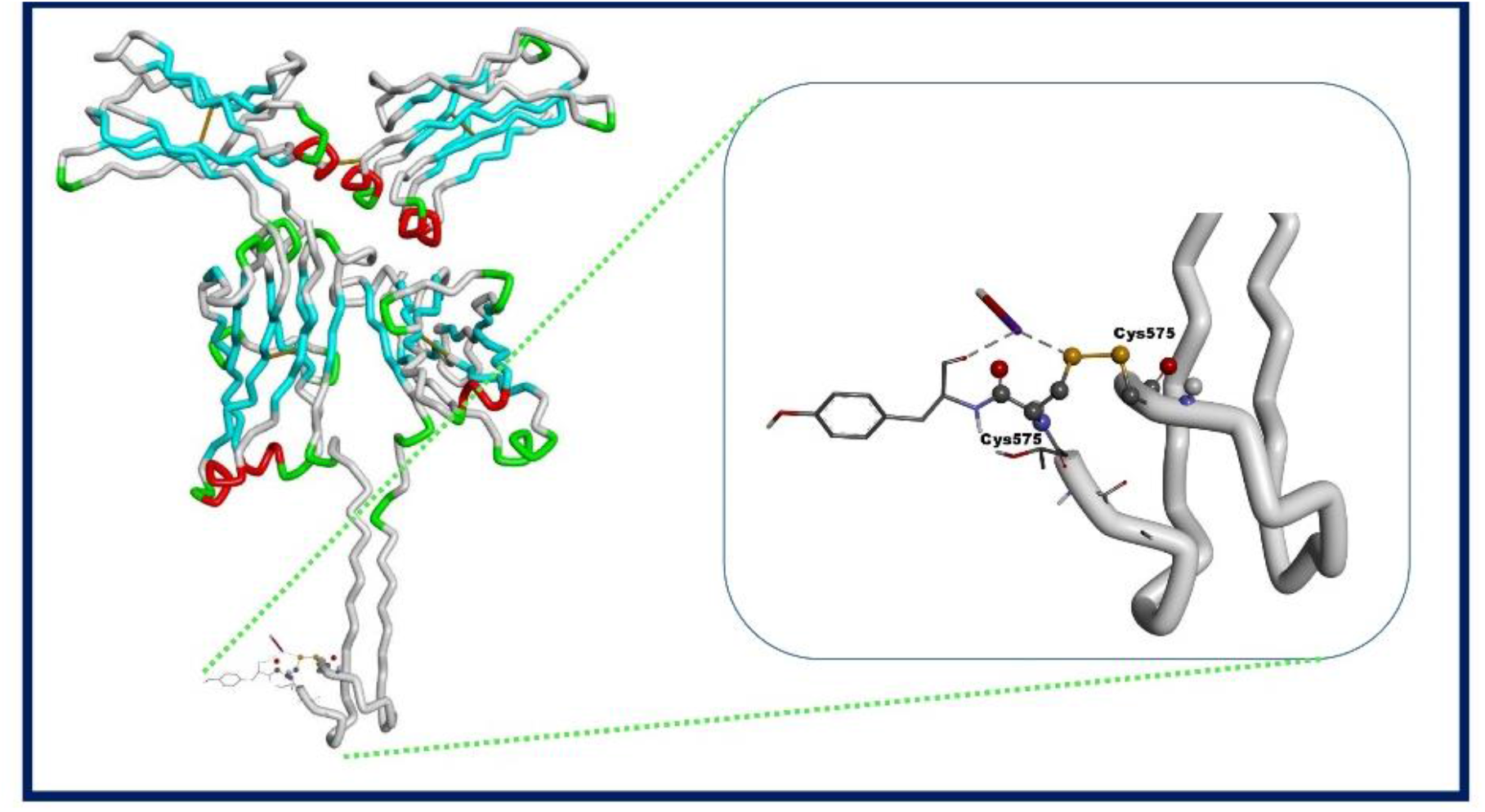
Potassium interaction in the disulfide bond formed by the sulfur atoms between the two cysteines at the 575th position in the IgM F and G chains.

## Discussion

IgA is an excess of Ig in the lumen secretion. Another Ig found in excess in this secretory fluid is IgM. IgM is a part of the innate defense system together with natural killers (NK), macrophages, dendritic and mast cells (19). Therefore, both IgM and IgA are important components of the immune system, which also plays a role in the first line of the defense system.

Igs can have different weights depending on their monomeric or polymeric structure. Naturally the heaviest Ig is IgM (900,000 Daltons) followed by IgA (320,000 Daltons). If there is no extreme property due to the laws of physics, the lighter one of similar molecules moves more easily. It is known that while Ig with monomeric structure moves by passive transport in their transport from the lamina propria to the apical surface in the respiratory system, pIgs move by active transport(18). In Ig transport from basolateral to apical surface, a molecular component such as SC for pIgs or molecular weight factor for energy requirement should also be considered.

In addition, Ig mobilization, including Ig rotations and changes in hinge angle, is very important for Ig to recognize and bind epitopes (21,22). The synthesis of different Ig fragments such as nanobody (single domain antibody), which has increased in the last 4 decades, has a very beneficial effect in the treatment of different diseases (23). In developing antibody treatment technologies, low weight Ig or Ig fragments have proven themselves in their therapeutic potential.

Because of the naturally higher number of Fab fragments on pIgs, pIgs can bind more antigens than monomeric Igs. However, if Fab fragments are not filled to full capacity in pIg’s, their avidity may decrease. Ag uptake in all Fab fragments of an Ig with high avidity such as IgM will greatly contribute to the formation of an early and strong immune response for the body.

While the IgA dimeric structure is held together by the J chain, both the J chain and intermonomeric disulfide bonds have an important place in the construction in the IgM pentamer. Disulfide bonds in the Ig structure have a very important role in keeping the structure stable. J chains also have many disulfide bonds. These disulfide bonds can be easily destroyed by nucleophilic attack via OH^−^ ions(10-12). Intermonomeric disulfide bonds, on the other hand, vary depending on the Ig amino acid sequence, but in our study, they were detected between the cysteine amino acids 414-414 and 575-575 in the IgM pentamer (The heavy chain amino acid sequence in the IgM F monomer is given in the supplement file).

In the context of our molecular docking simulations, we believe that it may be useful to give a brief explanation about the hypothesis established in this study: As it is known, KOH has the ability to react rapidly and aggressively with other molecules in its environment (both intracellular or extracellular) due to its molecular structure. This nucleophilic ability of KOH in organic chemistry is due to its behaviour as an OH^−^ source, and this highly nucleophilic OH^−^ ion has the ability to attack polar bonds in both inorganic and organic molecules. Thus, by nature, KOH is a polar molecule. S-S bonds in human Igs are strong, with an average bond dissociation energy of 60 kcal/mol (251 kJ mol^−1^). However, although these bonds are about 40% weaker than C-C or C-H bonds, disulfide bonds are considered “*weak bonds*” in many molecules. Also, reflecting the polarizability of the divalent sulfur atom, S-S bonds are susceptible to cleavage by polar chemicals (both electrophiles and nucleophiles) (24). Therefore, the possibility of KOH breakage of the S-S bonds in human Igs could be considered, and this process may contribute to the dissociation of these macromolecular structures consisting of multiple subunits (like holoenzymes) into their monomeric subunits.

The reactivity of KOH towards the Cys109-Cys134 S-S bond(-1.94 kcal/mol) in IgA is also important because in this region KOH directly attacks Cys134, the residue which contributes to the formation of the S-S bond (Figure 5).

Other interactions in the J chain in IgA were -2.52 kcal/mol and -2.12 kcal/mol (Cys13-Cys101 / Cys72-Cys92), respectively. The interactions took place with atoms near the disulfide bond. In this study, a single molecule of KOH was used. In our study, the interaction of disulfide bonds between IgM Cys414-Cys414 amino acids with KOH was formed by multiple KOH application. This makes us think that potassium-sulfur interactions may occur in a much larger number by increasing the number of KOH molecules or changing the location of the KOH molecule (Figure 5 and Figure 6). Since all of these interactions are exergonic reactions, it makes us think that S-S bonds can be destroyed.

Likewise, in the IgM J chain, the binding energies were -2.34 kcal/mol, -1.90 kcal/mol and - 2.07 kcal/mol(Cys12-Cys100 / Cys108-Cys133 / Cys14-Cys575), respectively. No interaction between potassium and sulfur was observed in these bindings. In silico study of a single KOH molecule between intermonomeric Cys414-Cys414, it was observed that the potassium atom did not bond to the sulfur atom. However, the presence of many KOH molecules in the environment created an interaction energy of -1.05 kcal/mol between the sulfur atom in the Cys414-Cys414 disulfide bond and the potassium atom. In the vicinity of this binding site, potassium formed an electrostatic bond with oxygen (416. amino acid (Aspartic acid) and 417. amino acid (aspartic acid) with oxygen atoms). The electrical charge of oxygen was -0.63 eV, while the electrical charge of sulfur was -.0.09 eV. This suggests that the sulfur atom in the disulfide bond interacts with potassium after the oxygen bond is saturated with the increase in potassium in the medium. This scenario seems more biologically appropriate. However, in the disulfide bond formed by the amino acids Cys575-Cys575 between the heavy chains (G and F heavy chains of IgM), -0.93 kcal/mol binding energies were present even in the presence of a single KOH molecule in the environment (**Table-1**) (**Figure 7**) (IgM heavy chain amino acid sequence is given in the supplemental file).

Since interchain disulfide bonds are more exposed to solvents than intrachain disulfide bonds, solvents affect interchain disulfide bonds more (25). These interactions may increase the potential to break the J chains in dimeric IgA and pentameric IgM, and the S-S bonds between monomers in IgM. It is thought that the IgM intermonomeric disulfide bonds that we targeted with KOH in our study are the most suitable bonds that can be targeted for interchain solvent interaction and are perhaps the most suitable bonds for the purpose of the study. These bonds are shown in the graphic image in 2D and in the video image in 3D (**Figure 8, Video 1**).

**Figure-8:**
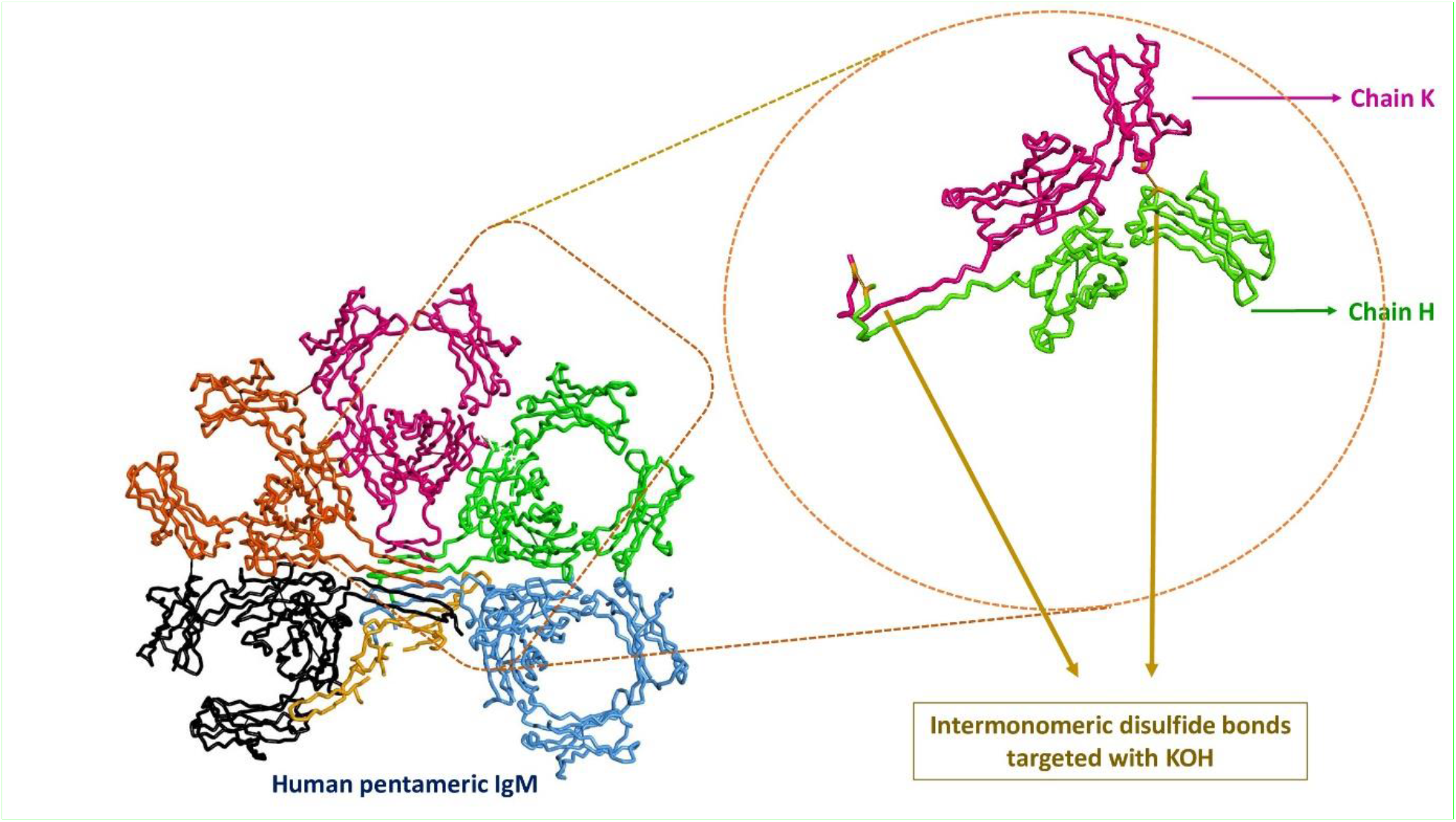
Disulfide bonds (between amino acids Cys414-Cys414 and Cys575-Cys575) in between two different monomers in IgM, whose interaction energies are calculated by targeting with KOH.

**Video-1:**
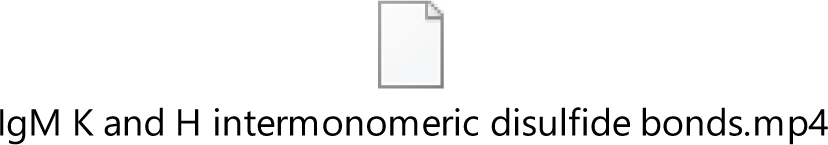
Video image of disulfide bonds between two different monomers in IgM, between amino acids Cys414-Cys414 and Cys575-Cys575, whose interaction energies are calculated by targeting with KOH. Green and pink colors represent single heavy chains of different monomers in IgM. The orange color represents the disulfide bonds connecting the two monomers.

In the light of this information, KOH with an alkaline molecular structure has the potential to monomerize pIg in the respiratory lumen.

Monomerization of pIgs will reduce the molecular weight of IgA and IgM. This feature can give IgA and IgM the ability to increase mobility. It will recognize monomeric IgA and IgM epitopes with increased mobility and will be able to bind more Ag structures. The number of Ag bound by individual monomers may be greater than the Ag number of a single pIg. This can increase both affinity and avidity of pIgs.

Monomeric IgA and IgMs can easily pass from the apical surface to the basolateral area by passive transport without the need for SC with their antigen load. This will lead to early and strong immune response with early recognition of Ag.

KOH molecules can reach the lungs by inhalation. In the evaluation of the studies, it was discussed that the application of the KOH molecule by inhalation in a solution compatible with the interstitial serum(0.89% NaCl in distilled water and pH:8.9) does not cause toxic effects in the lungs, and also has a mucolytic and alkalinizing effect in the mucus (26).

The KOH molecule reaching the lungs as a result of inhalation of KOH can interact with Ig disulfide bonds in the mucus. Even if there is no disease in the lungs, KOH and IgM monomers can form. With the passage of monomerized IgM fragments into the systemic circulation without any Ag load, IgM monomers can help to capture Ag structures in diseased areas in all distant organs, especially in infections, cancer and autoimmune diseases, and present them to the immune system. This may cause early opsonization in the diseased area for diseases in the body. This may have a positive effect on increasing the effectiveness of chemotherapeutic agents used in patients and increasing the innate immune response and ensuring the continuity of body resistance.

More studies are needed on this subject, as potential breaks in disulfide bonds or bitterness changes in the hinge regions rich in disulfide bonds and between the heavy and light chain may lead to different immune responses.

## Conclusion

Intermolecular exergonic reactions have developed between KOH and S-S bonds in the J chain of pIgs.

Intermolecular exergonic reactions have developed between intermonomeric S-S bonds in KOH and IgM pentamer.

In case of disease in the body, there may be a potential for early and strong initiation of immune response in all tissues by the monomerization effect of KOH in pIg.

## Supporting information

Graphical abstract and others supplemental figures

## Acknowledgment

I would like to thank Ph.D. Erman Salih İstifli for his help in the simulation studies.

