## Supplementary material for "For an early and strong immune response in the monomerization of polymeric immunoglobulins intermonomeric interaction relationship of potassium hydroxide": Graphical abstract and others supplemental figures

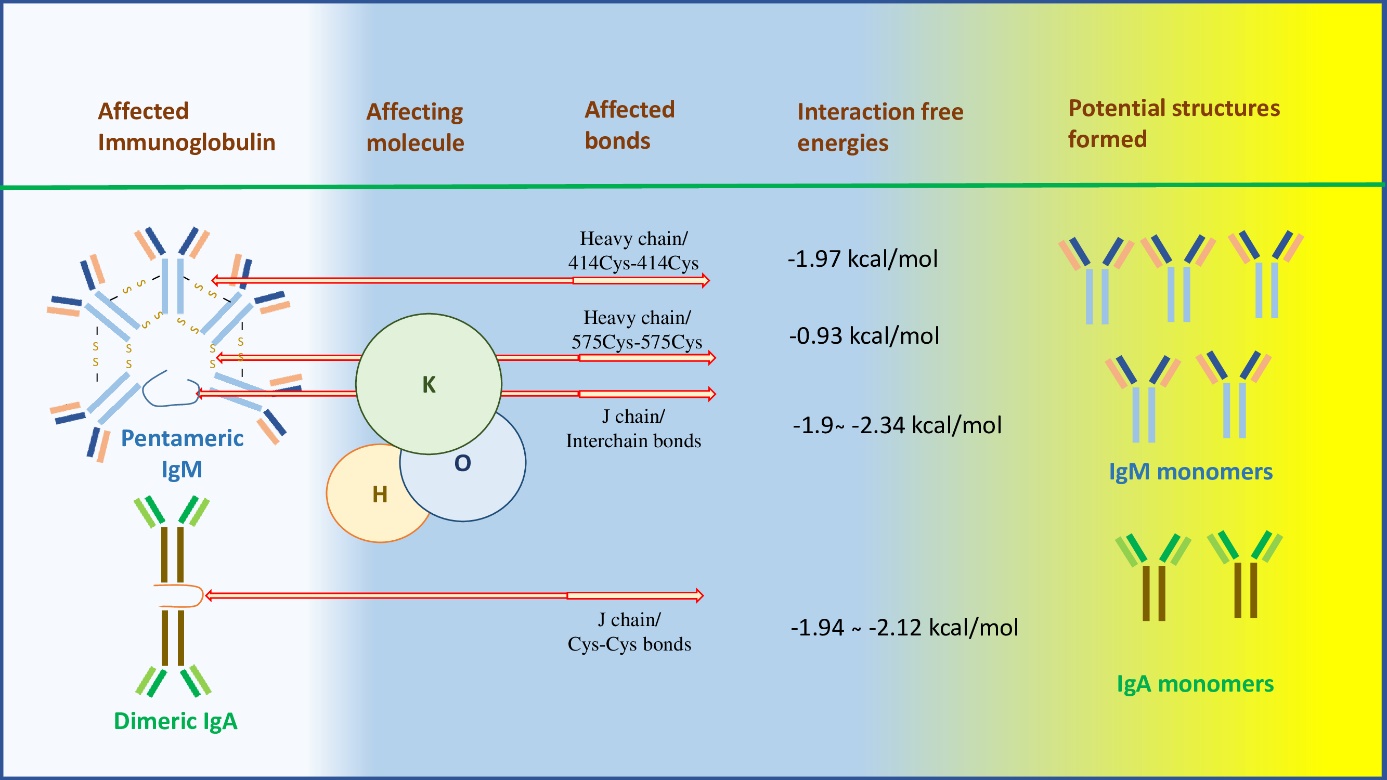

**Graphical abstract:** Graphic with abstract of the study.

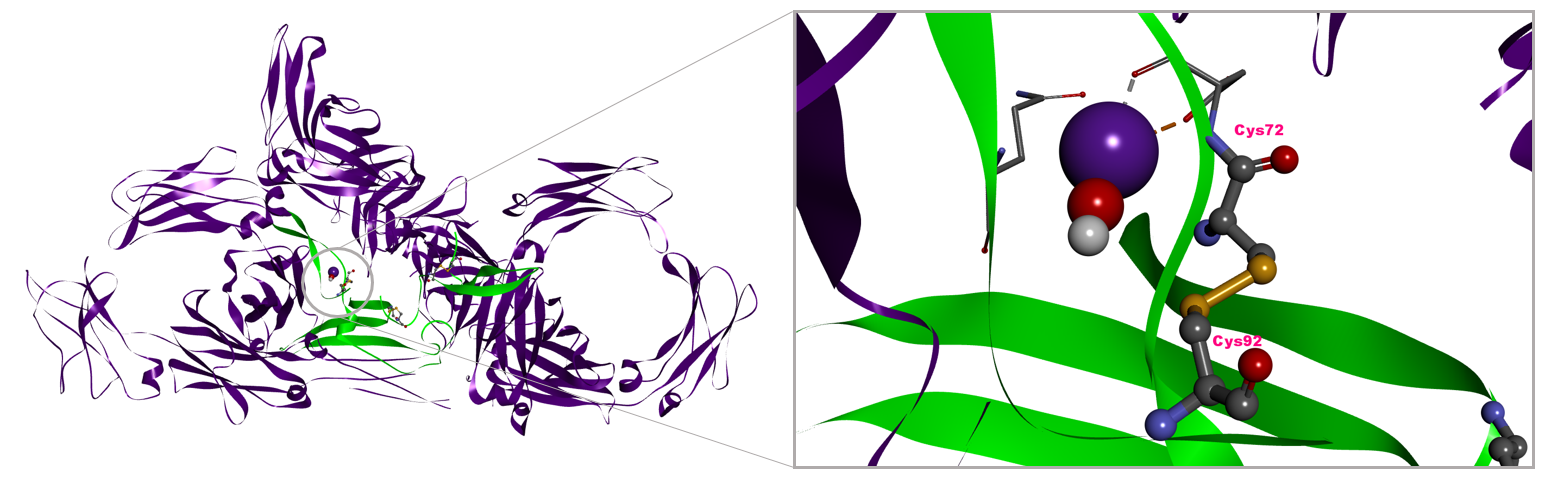

**Figure 1.** Top-ranked conformation of KOH in complex with the human dimeric sIgA in the vicinity of Cys72-Cys92 disulfide bond. The orange dashed line represent electrostatic, whereas the grey dashed line represent metal-acceptor interactions, respectively. Image rendering was performed with DS Studio v16 software.

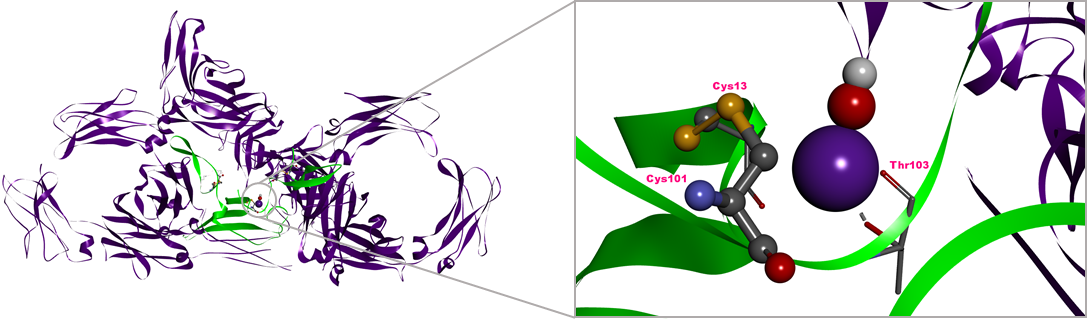

**Figure 2**. Top-ranked conformation of KOH in complex with the human secretory dimeric immunoglobulin A (sIgA) in the vicinity of Cys13-Cys101 disulfide bond. The single grey dashed line represent metal-acceptor interactions, respectively. Image rendering was performed with DS Studio v16 software.

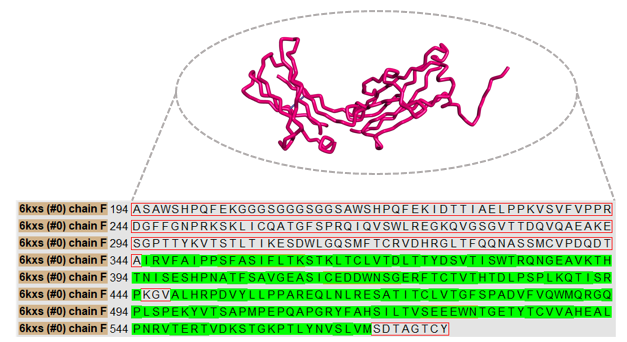

**Figure 3.** Amino acid sequence of the F chain (immunoglobulin heavy constant µ) of the human immunoglobulin pentameric M antibody (PDB ID: 6KXS). Missing residues in the crystal structure are surrounded with a red line, whereas the retained residues in the crystal structure are depicted with green background. The amino acid sequence image was created with the Chimera 1.16r program.

In the supplement video file

**Video-1:** Video image of disulfide bonds between two different monomers in IgM, between amino acids Cys414-Cys414 and Cys575-Cys575, whose interaction energies are calculated by targeting with KOH. Green and pink colors represent single heavy chains of different monomers in IgM. The orange color represents the disulfide bonds connecting the two monomers.
